# Loss of GluN2A Selectively Impairs Excitatory Synaptic Integration in Hippocampal Somatostatin Interneurons

**DOI:** 10.64898/2026.11.11.763692

**Authors:** Daniel W Hunter, Hannah Kaese, Peter C Kind, David JA Wyllie

## Abstract

N-methyl-D-aspartate receptors (NMDARs) are widely considered key determinants of excitatory synaptic kinetics and activity-dependent plasticity, yet their specific contribution to postsynaptic potentials across distinct hippocampal CA1 cell types remains poorly defined. Null variants in *GRIN2A* are strongly associated with epilepsy-aphasia spectrum disorders, and GluN2A loss has recently been linked to delayed electrophysiological maturation of parvalbumin interneurons, a process regulated by somatostatin interneuron activity. Here, we used *Grin2a* haploinsufficient and knockout rats to determine how GluN2A depletion affects excitatory synaptic transmission, integration and firing across CA1 pyramidal neurons, parvalbumin interneurons and somatostatin interneurons. Using whole-cell patch-clamp recordings in acute hippocampal slices, we found that NMDARs made no discernible contribution to shaping Schaffer collateral-evoked EPSP kinetics, temporal summation or spike recruitment in CA1 pyramidal neurons at physiological resting potentials. Hebbian long-term potentiation at pyramidal neuron Schaffer collateral synapses was also preserved. Parvalbumin interneuron EPSP kinetics and synaptically evoked firing were similarly unaffected. By contrast, GluN2A loss selectively prolonged EPSP decay in CA1 somatostatin interneurons, increasing EPSP area, enhancing frequency-dependent temporal summation and elevating action potential output during repetitive and theta-frequency synaptic stimulation. These changes occurred alongside an increase in intrinsic excitability, indicating a cell-type-specific alteration by GluN2A loss. Selective enhancement of somatostatin interneuron recruitment provides a candidate mechanism - upstream of delayed parvalbumin interneuron maturation - and may contribute to hippocampal network dysfunction in *GRIN2A*-related disorders.

## Introduction

Healthy brain function is contingent on a finely tuned balance between excitatory drive, which encodes and propagates information, and inhibitory control, which constrains, shapes and coordinates that signalling through precisely organised patterns of connectivity, timing and plasticity. Synaptic N-methyl-D-aspartate receptors (NMDARs) are central to excitatory neurotransmission and activity-dependent plasticity. Their distinctive biophysical properties - voltage-dependent Mg²⁺ block, high Ca²⁺ permeability, and slow gating kinetics - position them as coincidence detectors at excitatory synapses, integrating pre- and postsynaptic activity, gating intracellular signalling cascades governing synaptic strength (Collingridge, 2025). NMDARs are obligate heterotetramers, comprising two GluN1 subunits combined with GluN2 subunits (GluN2A-D) that confer distinct kinetic properties on the assembled receptor. In the adult hippocampus, GluN2A and GluN2B are the predominant GluN2 subunits, with the GluN2A subunit conferring faster deactivation kinetics relative to GluN2B (Paoletti et al., 2013; Wyllie et al., 2013). The developmental regulation of GluN2A and GluN2B expression is well established throughout the brain, and the relative localisation of these subunits at mature synapses has long been proposed to influence the threshold and direction of synaptic plasticity (Keith et al., 2024; Kellermayer et al., 2018).

The GluN2A subunit has attracted particular interest in the context of hippocampal long-term potentiation (LTP). Early work suggested opposing roles for GluN2A and GluN2B in LTP induction, with GluN2A proposed to favour potentiation and GluN2B supporting synaptic depression (Liu et al., 2004; Massey et al., 2004). However, this simplistic model has not survived more robust examination: pharmacological and genetic deletion of GluN2 subunits produces highly variable effects on LTP and LTD magnitude, largely depending on the specific induction protocol and preparation (Berberich et al., 2007; Shipton and Paulsen, 2014). The consensus view now favours a more nuanced role for GluN2A, in which subunit composition shapes the kinetics of synaptic Ca²⁺ signals and the downstream engagement of plasticity-related signalling cascades rather than acting as a binary gate on LTP (Shipton and Paulsen, 2014). The consequences of GluN2A loss for synaptic physiology in the hippocampus therefore remain incompletely resolved, particularly at the level of intact circuit function where metaplastic processes and local dendritic inhibitory drive add yet further complexity.

The GluN2A subunit is encoded by *GRIN2A* in humans. Loss-of-function *GRIN2A* variants - including missense, truncating and null mutations - are among the most frequent monogenic causes of epilepsy-aphasia spectrum disorders encompassing self-limited epilepsy with centrotemporal spikes, Landau-Kleffner syndrome and epileptic encephalopathy with continuous spike-and-wave during sleep (CSWS) (Lesca et al., 2013; Strehlow et al., 2019). Despite this, the cellular and circuit-level mechanisms through which partial or complete loss of GluN2A drives pathological network excitability remain poorly understood. A fundamental unresolved question is whether the consequences of GluN2A loss are distributed homogeneously across neuronal populations, or instead concentrated in specific, selectively vulnerable subpopulations. Interneuron dysfunction is increasingly recognised as a convergent mechanism across many genetic epilepsies, and disruption of parvalbumin and somatostatin interneuron activity in particular has been proposed to contribute to hippocampal network instability (Magloire et al., 2019; Sessolo et al., 2015; Wengert et al., 2021; Yuan et al., 2024). Understanding whether GluN2A loss differentially impacts inhibitory interneuron populations compared to pyramidal cells would therefore provide a mechanistic framework relevant to the pathophysiology of *GRIN2A*-related disorders.

GABAergic interneurons express functional NMDARs and receive glutamatergic input from both local projections and afferent pathways, yet their NMDAR subunit composition and the functional consequences of subunit loss have received comparatively little attention (Moreau and Kullmann, 2013). Distinct inhibitory subtypes - defined by molecular markers, morphology, and connectivity - occupy specific positions in hippocampal circuits and perform highly specialised functions (Booker and Vida, 2018). Parvalbumin-expressing interneurons target the perisomatic region of pyramidal cells and provide fast, powerful feedforward inhibition that controls spike timing and supports gamma oscillations (Hu et al., 2014). Somatostatin-expressing interneurons, particularly those of the oriens-lacunosum moleculare (OLM), occupy a distinct functional niche: they receive excitatory input from local CA1 pyramidal cells and project their axons to the distal dendrites of these pyramidal cells in stratum lacunosum-moleculare, where they selectively inhibit entorhinal cortical input while disinhibiting CA3-driven signals through a parvalbumin interneuron disinhibitory pathway (Chamberland et al., 2023; Honoré et al., 2021). Somatostatin cells are thus well placed to gate the relative influence of cortical and subcortical inputs into the CA1 (Leão et al., 2012).

The subunit composition of NMDARs at excitatory synapses onto these two interneuron classes, and the functional consequences of GluN2A loss for their activity, are not known. This question is made more pressing by recent evidence that GluN2A loss produces a transient delay in the electrophysiological maturation of hippocampal parvalbumin interneurons (Camp et al., 2023). Parvalbumin interneuron maturation is known to depend on somatostatin interneuron activity during a critical developmental window (Mòdol et al., 2024). It has also been demonstrated that excitatory synapses onto somatostatin interneurons compensate for the loss of GluN2A through an increased reliance on GluN2B-containing receptors, as shown by an increased fraction of ifenprodil block in the *Grin2a*^-/-^ rat, relative to wildtype (Booker et al., 2021). We therefore hypothesised that somatostatin interneurons may be selectively sensitive to *GRIN2A* loss, such that reduced GluN2A expression - and compensation by GluN2B - would disproportionately modulate their functional integration relative to other hippocampal cell populations.

In the present study, we employ whole-cell patch clamp electrophysiology in acute hippocampal slices from a *Grin2a* heterozygous and homozygous knockout rat model to systematically examine the cell-type-specific consequences of GluN2A loss on synaptic physiology, intrinsic membrane properties, temporal summation and activity-dependent firing across CA1 pyramidal neurons, parvalbumin interneurons and somatostatin interneurons. We demonstrate that GluN2A loss does not alter synaptic physiology, temporal integration or Hebbian LTP at the Schaffer collateral pyramidal cell synapse, and leaves parvalbumin interneuron function largely intact. In contrast, somatostatin interneurons exhibit a selective, gene-dose-dependent slowing of EPSP kinetics, markedly enhanced temporal summation of synaptic inputs, reduced dendritic complexity and elevated firing across stimulus frequencies including the physiological theta band. These findings identify somatostatin interneurons as a primary cellular locus of GluN2A depletion in hippocampal CA1 microcircuits and suggest that their selective hyperactivation may constitute a key mechanism linking *GRIN2A* haploinsufficiency to network dysregulation and pathophysiology.

## Methods

### Animals

A *Grin2a* knockout Long-Evans Hooded rat line was used in this study, in which the exon of the *Grin2a* gene which encodes key pore forming domains of GluN2A and contains N615 was deleted, resulting in a non-translated gene product. Depending on the transcript ID and genome assembly the exon number varies, and in previous publications this exon has either been named as exon 8 (Strehlow et al., 2019) or exon 11 (Yasmin et al., 2025). These rats were generated by pronuclear microinjection of mRNA encoding the enzyme Cas9, single guide RNA binding to 5′ and 3′ of exon 11 of *Grin2a*, and a pUC57 plasmid containing 2142 base pairs of donor DNA into single-cell Long-Evans Hooded rat embryos before being implanted into pseudo-pregnant mothers. The resulting live births were screened by PCR for either insertion of donor DNA via homology directed repair or genomic deletions due to repair by non-homologous end joining of double stranded. Rats where non-homologous end joining had occurred showed a large 1065 base pair exonic deletion. The single guide RNA design and generation of the founder animals was performed by Horizon Discovery Group plc (St. Louis, MO, USA). Confirmation by Sanger sequencing was performed by PCR amplification of a genomic region spanning the donor DNA section and including endogenous DNA to 5′ and 3′.

Animals heterozygous for the deletion were then crossed to give offspring that were either wild-type (WT, *Grin2a*^+/+^), heterozygous (HET, *Grin2a*^+/-^) or homozygous (KO, *Grin2a*^-/-^). Offspring, both male and female, from these litters were culled for tissue between postnatal days 21 and 34 (P21-34). For experiments in which it was necessary to visualise inhibitory interneurons, we crossed *Grin2a*^-/-^ animals with a VGAT-mVenus reporter line. In this reporter, a bacterial artificial chromosome in which the fluorescent protein Venus is expressed under the vesicular GABA transporter (VGAT) promotor, has been randomly intercalated into the genome permitting identification of GABAergic interneurons in the mature and developing rat brain. This cross produced VGAT-mVenus+ *Grin2a*^+/-^ offspring which could then be crossed with *Grin2a*^+/-^ animals to produce *Grin2a* WT, HET and KO pups expressing mVenus in inhibitory interneurons. Rats were housed under a standard 12:12 h light/dark cycle and received food and water ad libitum. Experiments performed in this study were in accordance with the United Kingdom Animals (Scientific Procedures) Act of 1986 under the Home Office licence PP2262369.

### Acute hippocampal slice preparation

Animals were decapitated under isoflurane anaesthesia, and the brains quickly removed and submerged in carbogenated (95% oxygen, 5% CO2) ice-cold dissection buffer containing the following (in mM): 86 NaCl, 1.2 NaH_2_PO_4_, 2.5 KCl, 25 NaHCO_3_, 25 glucose, 75 sucrose, 0.5 CaCl_2_ and 7 MgCl_2_. Four hundred micrometre horizontal hippocampal slices were prepared using a vibratome (Leica VT 1200S). Slices were collected in dissection buffer and incubated for 30 minutes at 35°C after which slices were maintained at room temperature until recording.

### Whole-cell patch-clamp recordings

For electrophysiological recordings, slices were placed in a submersion chamber heated to 34°C with an inline heater (LinLab Scientifica) and perfused with carbogenated artificial cerebrospinal fluid (aCSF) at a rate of 6 mL/min, containing the following (in mM): 124 NaCl, 1.2 NaH_2_PO_4_, 2.5 KCl, 25 NaHCO_3_, 20 glucose, 2 CaCl_2_ and 1 MgCl_2_, bubbled with 95% oxygen and 5% CO_2_. Slices were visualized under infrared differential inference contrast microscopy with a digital camera (SciCamPro, Scientifica) mounted on an upright microscope (model BX61-WI, Olympus) with a 60× water-immersion objective lens [1.0 numerical aperture (NA), Olympus]. For patch clamp experiments in inhibitory interneurons, fluorescently labelled cells were targeted for whole-cell patch clamp under combined infrared differential interference contrast and epifluorescence illumination. Recorded cells were filled with biocytin via the patch pipette to permit post-hoc morphological reconstruction and verification of cell identity. parvalbumin interneurons were identified on the basis of soma location in stratum pyramidale, a fast-spiking discharge pattern characterised by high-frequency action potential firing with minimal adaptation, and confirmed by post-hoc biocytin reconstruction. somatostatin interneurons were identified by soma localisation within stratum oriens, consistent with the OLM cell subtype and confirmed by post-hoc morphological reconstruction. The characteristic somatostatin cell morphology, with a horizontally oriented dendritic arbour confined to stratum oriens, was evident in reconstructed somatostatin interneurons across all genotypes. Where Schaffer collateral stimulation was required by the experimental protocol, a parallel bipolar electrode (FHC) was placed in the stratum radiatum between the CA3 and CA1 and stimuli were delivered via constant current isolated stimulator (Digitimer). Whole-cell recording electrodes were pulled using borosilicate glass capillaries (1.5mm outer, 0.7mm inner diameter) on a horizontal puller (P-97, Sutter Instrument Co.). For current-clamp recordings, pipettes were filled with a potassium gluconate-based solution (in mM: 120 potassium gluconate, 20 KCl, 4 NaCl, 0.1 EGTA, 10 HEPES, 4 Na_2_-ATP, 0.3 Na_3_-GTP, 10 Na_2_-phosphocreatinine, pH 7.3, 290-310mosmol l^-1^), with a final resistance of 3-5 MΩ. For voltage-clamp recordings, a caesium gluconate-based internal solution (in mM: 140 Cs-Gluconate, 3 CsCl, 0.2 EGTA, 10 HEPES, 2 Mg-ATP, 2 Na_2_-ATP and 0.3 Na_3_-GTP, 5 QX314Cl, pH 7.35, 290-310 mosmol l-1), with a final resistance of 3-5 MΩ was used. Biotinylated-lysine (0.1%; Biocytin, Invitrogen, UK) was included in recordings of identified interneurons where dendritic reconstructions were required. All recordings were performed using a Multiclamp 700B amplifier (Molecular Devices). Voltage and current signals were low-pass filtered at 10 and 2 kHz, respectively, using the built in Bessel filter of the amplifier, digitized at 20 kHz (Digidata 1440, Molecular Devices) and acquired with pCLAMP software (pClamp 11, Molecular Devices). Data were analysed offline using ClampFit software (Molecular Devices) and custom-made MATLAB scripts. Series resistance (R_S_) was constantly monitored throughout the recordings but was not compensated. Recordings were abandoned if initial R_S_ exceeded 30 MΩ or changed by more than 20% over the time course of the recording, if I_hold_ was less than −200pA in voltage-clamp recordings at −70mV or resting membrane potential was more depolarized than −50mV in current-clamp recordings.

For evoked excitatory postsynaptic current (eEPSC) recordings following break-in to whole cell configuration, the amplitude level was adjusted on the constant current stimulator to evoke and eEPSC of approximately 200pA. When a stable response was achieved, 30 AMPAR-eEPSCs were recorded were at 3s intervals. For AMPAR/NMDAR composite eEPSCs, the cell was subsequently held at +40 mV to unblock the NMDAR component. All composite eEPSCs were recorded at +40 mV using the same stimulation intensity as for AMPAR-eEPSCs. For experiments conducted at +40mV, 5_μM_ gabazine was supplemented to the aCSF to isolate excitatory postsynaptic currents and potentials. All eEPSC amplitudes were recorded at the peak of the eEPSC. Halfwidth was determined as the time duration of a synaptic current waveform measured exactly at half of its maximum peak amplitude. Decay time constants were calculated from a standard exponential curve (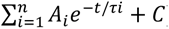) fit to the decaying phase of the eEPSC. For all kinetic properties, measurements were performed on averaged synaptic events, to minimise measurement and fitting artifacts. NMDAR/AMPAR eEPSC amplitude ratios were calculated from the −70mV AMPAR-eEPSC peak amplitude divided by the +40mV composite eEPSC amplitude measured 50ms after the peak. For evoked excitatory postsynaptic potential (eEPSP) recordings, the amplitude level was adjusted on the constant current stimulator to evoke and eEPSP of approximately 5-10mV. When a stable response was achieved, 30 eEPSPs were recorded were at 3s intervals, while injecting a biased current to maintain the resting potential at −70mV. For NMDAR-unmasked eEPSPs, the biased current was increased to maintain the membrane potential at +40 mV. All composite eEPSPs were recorded using the same stimulation intensity as for −70mV eEPSPs. For quantification, eEPSP amplitude, halfwidth and decay time constants were calculated from average synaptic events, as described above.

Synaptic summation of eEPSPs was measured with trains of 5 stimuli delivered at 100, 80, 60, 40 and 20Hz (10 s inter-sweep interval); 16 traces were collected for each frequency tested. Stimulus intensity was adjusted to elicit a first subthreshold eEPSP of approximately 5-10mV in amplitude and was kept constant throughout the stimulus trains. For experiments in which action potential firing was prevented, to allow visualisation of summating suprathreshold eEPSPs, 5mM QX-314 was included in the pipette solution. Spike latency for a given pulse train was determined as the minimum time duration between first stimulus onset and action potential threshold, where threshold was defined as the point at which *∂V*/*∂t* > 20*mVms*^−1^. Evoked EPSP area was calculated by measuring the area between the EPSP waveform and the baseline (∫(*V_m_*(*t*) − *V_baseline_*)*∂t*), reflecting the total charge transferred across the membrane during the synaptic potential. For analysis, the determined eEPSP area in response to a given stimulus was normalised to the first response in the pulse train. Spike count was used for action potential output in hippocampal interneurons in place of spike latency, as these cells displayed capacity to fire repetitively in response to broad rage frequency inputs. Spike count was determined as the average number of action potentials detected across the pulse train.

For LTP recordings, the membrane potential CA1 pyramidal neurons was voltage clamped at −70mV. Then the constant current stimulator was adjusted to elicit eEPSCs with a peak amplitude of approximately 200pA. Baseline eEPSCs were recorded at 0.1Hz. The baseline stability criterion was defined as less than 10% change in eEPSC peak amplitude, calculated from a linear fit of baseline period. If stability was not achieved within 10 min of breakthrough, the recording was abandoned. Following 6 min of stable recording, LTP was induced, with 2 × 100 Hz of tetanus stimuli, at the same strength as the baseline eEPSC. Following induction, 30 min of 0.1Hz eEPSCs were recorded to assess potentiation of synaptic response, quantified as the eEPSC peak amplitude relative to baseline average.

For spontaneous excitatory and inhibitory postsynaptic current (sEPSC and sIPSC, respectively) recordings, CA1 somatostatin interneurons were visually identified by horizontal somatic morphology located within the stratum oriens. Upon break-in to whole cell configuration, cells were held at a membrane potential of −70mV, which is close to the reversal potential for chloride, to reveal isolated sEPSCs. To isolate inhibitory synaptic events, the holding potential was shifted to 0mV, close to the reversal potential for AMPA- and NMDA receptor mediated currents, to record GABA_A_ receptor-mediated sIPSCs. Spontaneous synaptic activity was recorded for 6 minutes in each configuration. Prior to event detection, recordings were processed with the electrical interference filter in ClampFit (Molecular Devices), to reject 50Hz line frequencies and their harmonics. Events were then detected using the template search mode, only events with an amplitude greater than 6pA were included for analysis. Synaptic frequency measures were averaged per cell and then per animal. Quantification of peak amplitude and decay time constant (τ) was calculated as described above. Kinetic analysis was performed by fitting to an average of all event detections from a given cell. This approach was used since single-event values were often unreliable due to noise or nearly linear decay. Cell averages were then combined to generate an animal average for statistical investigation.

Intrinsic physiology of inhibitory interneurons was recorded in current-clamp mode, administering a series of 500ms current injection steps from −100 to +400pA, in 25pA increments. Input-output curves were plotted from the average action potential frequency per current step. Resting membrane potential was determined from the inter-step interval, after steady state potential had been reached. Rheobase was determined as the current injection step in which an action potential was first elicited. Membrane time constant and input resistance were was calculated from an average of 30 hyperpolarising current steps (−20pA, 1s), where membrane time constant was calculated from a standard exponential curve fit to the initial hyperpolarising phase of the voltage response. Input resistance was calculated from the steady-state voltage at the end of the current step, by Ohms law (*R_in_* = *∂V*⁄*∂I*).

Activity-dependent firing of CA1 somatostatin interneurons was assessed using whole-cell current-clamp recordings. somatostatin interneurons were identified and patch clamped as described above, and recordings were performed in current-clamp mode without bias current injection. Summating subthreshold eEPSPs were elicited at 8Hz for 6 s, followed by a 6 s inter-train interval. This stimulation protocol was repeated 10 times for each recorded cell. Membrane voltage responses were recorded throughout each train, and action potentials were detected using a derivative-based detection criterion (*∂V*/*∂t* > 20*mVms*^−1^). For each train, the total number of action potentials was quantified as a measure of stimulus-evoked firing output. Spike-frequency adaptation was assessed using an adaptation index, this was calculated as the number of action potentials evoked during the final portion of the train divided by the number evoked during the initial portion of the train, such that lower values indicate greater adaptation across the stimulus train. The depolarisation envelope was quantified as the slow underlying membrane depolarisation generated during the stimulus train, measured as the 2ms immediately prior to stimulus onset of the final 8 stimuli, and expressed relative to the pre-train baseline. This measure was used to capture the cumulative depolarising drive and temporal summation produced by repeated theta-frequency synaptic stimulation.

### Immunohistochemistry and dendritic analysis

Morphological analysis of the recorded CA1 pyramidal cells and inhibitory interneurons was performed following whole-cell recording. Slices were fixed immediately after recording with 4% paraformaldehyde in 0.1 M phosphate buffer, overnight at 4°C. Slices were then rinsed three times for 15 minutes in phosphate buffered saline (0.1 M phosphate buffer + 0.9% NaCl; PBS). Slices were permeabilised with 0.3% Triton-X100 in PBS for 1 hour at room temperature, then incubated with Streptavidin conjugated to Alexa Fluor 633 (1:500, Invitrogen, Dunfermline, UK) diluted in 0.1% Triton-X100 in PBS, overnight at 4°C. Slices were rinsed as before in PBS and incubated with 2µM Hoechst 33342 in water for 15 minutes. Slices were washed in in PBS three times before one final wash in distilled water before mounting with fluorescence-protecting mounting medium (Fluoromount-G). Recorded neurons were visualized on a laser scanning confocal microscope (Zeiss LSM800 with AiryScan) with a 20× (N.A 0.75) objective lens for reconstruction. CA1 interneurons were 3-dimensionally reconstructed by combining multiple image z-stacks (1_μm_ steps) using the Simple Neurite Tracer plug-in for FIJI. Sholl analysis was performed on the 3-dimensionally segmented neurons, with a vector drawn from the centre of the soma to the greatest dendritic extent. Based on this distance, concentric shells (1_μm_ radius increments) were drawn from the origin (soma) and the number of dendritic intersections measured for each increment.

### Statistical analysis

Values are reported as mean ± SEM. Statistical comparisons were made using ‘N’ as number of animals. Data were averaged for each animal used, thereby reducing the risk of type-1 statistical errors resulting from pseudo-replication of the data. Average data throughout are presented as the average of each animal. Multiple group comparisons were carried out using ANOVA followed by post hoc pairwise tests, performed if the F test was significant for a main effect. Correction for multiple comparisons is made using Holm-Sidak method. P < 0.05 was considered statistically significant (∗P < 0.05, ∗∗P < 0.01 and ∗∗∗P < 0.001). Data were analysed using Prism (GraphPad Software, La Jolla, CA, USA).

## Results

### GluN2A loss does not alter pyramidal neuron synaptic physiology

Analysis of eEPSC decay at −70mV revealed no significant difference between WT, HET and KO tissue, consistent with predominantly AMPAR-mediated currents under near-complete Mg²⁺ block. Depolarisation to +40mV prolonged eEPSC decay in all genotypes and revealed a significant genotype effect, with decay tau increased in HET and further prolonged in KO relative to WT (Figure 1A-B; [F (2, 25) = 12.66 P=0.0002]). This gene-dose-dependent slowing is consistent with a shift toward slower GluN2B-containing NMDARs following loss of GluN2A. D-APV restored decay kinetics to values comparable to −70mV and abolished genotype differences, confirming that the effect was NMDAR-mediated.

**Figure 1.**
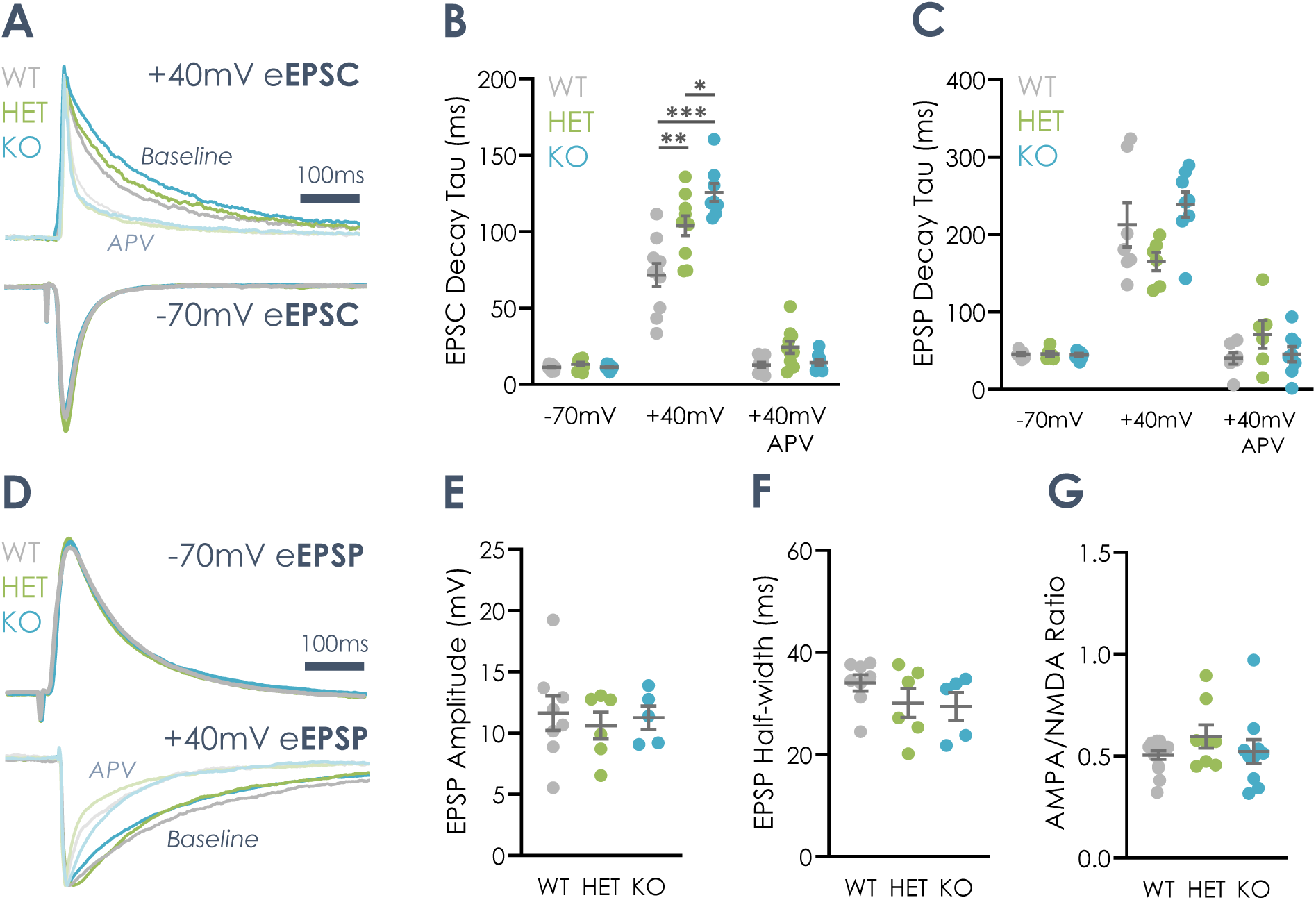
**Loss of GluN2A slows NMDAR-mediated eEPSC decay but leaves basal synaptic physiology of CA1 pyramidal neurons unaltered**. (A) Example eEPSC traces from a CA1 pyramidal neuron, recorded in voltage clamp at −70 mV, at +40 mV, and at +40 mV after wash-in of D-APV. Responses were evoked by stimulation of the Schaffer collaterals with a bipolar electrode. (B) eEPSC decay time constant (τ) in WT, HET and KO tissue at −70 mV, at +40 mV and at +40 mV with D-APV. At −70 mV, decay τ did not differ between genotypes. At +40 mV, decay was significantly slower in all genotypes, and there was a significant effect of genotype, with *Grin2a* loss slowing eEPSC decay. D-APV returned the +40 mV decay τ to −70 mV levels and abolished the genotype effect, confirming that the slowing is NMDAR-mediated (animal averages; N = 10 WT, 10 HET, 8 KO animals). (C) eEPSP decay τ for the same conditions as (B). At −70 mV, decay τ did not differ between genotypes. At +40 mV, decay was significantly slower, but there was no effect of genotype. D-APV returned eEPSP decay to −70 mV levels, again with no genotype effect (animal averages; N = 7 WT, 6 HET, 8 KO animals). (D) Example eEPSP traces from a CA1 pyramidal neuron, recorded in current clamp at −70 mV, at +40 mV, and at +40 mV with D-APV. (E) eEPSP amplitude at −70 mV did not differ between genotypes (animal averages; N = 8 WT, 6 HET, 5 KO animals). (F) eEPSP halfwidth at −70 mV did not differ between genotypes (animal averages; N = 8 WT, 6 HET, 5 KO animals). (G) AMPAR/NMDAR ratio did not differ between genotypes (animal averages; N = 14 WT, 8 HET, 10 KO animals). Data are mean ± SEM, with each point representing one animal. Statistical comparisons used ANOVA with Holm-Sidak post hoc tests.

In current clamp, eEPSP decay tau did not differ between genotypes at −70mV. Depolarisation to +40mV prolonged eEPSPs across genotypes, but no genotype-dependent difference was detected (Figure 1C-D; [F (2, 18) = 1.392 P=0.2742]). D-APV returned eEPSP decay to −70mV-like values in all genotypes. Thus, although GluN2A loss slows isolated NMDAR-mediated current deactivation under voltage clamp, this does not measurably alter the composite EPSP waveform in CA1 pyramidal neurons.

Additional measures of basal excitatory transmission were also unchanged. eEPSP amplitude, eEPSP halfwidth, and AMPA/NMDAR ratio did not differ significantly between WT, HET and KO tissue (Figure 1E-G; [eEPSP Amplitude: F (2, 16) = 0.1726, P=0.8430; eEPSP Halfwidth: F (2, 16) = 1.283, P=0.3041; AMPA/NMDA Ratio: F (2, 29) = 1.131, P=0.3366]). Together, these data indicate that loss of GluN2A alters NMDAR kinetics without substantially disrupting basal excitatory synaptic physiology at CA1 pyramidal neuron synapses.

We next tested whether altered NMDAR kinetics affected synaptic integration during repetitive Schaffer collateral stimulation. Spike latency during 100, 80, 60, 40 and 20Hz trains did not differ between genotypes (Figure 2B; [F (2, 18) = 0.6126, P=0.5528]). These findings are in agreement with previous work from the lab, showing no impact of *Grin2a* deletion on pyramidal cell summation (Yasmin et al., 2025). We further examined the impact of NMDAR blockade on pyramidal cell summation, revealing that D-APV produced no genotype-dependent change in spike latency (Figure 2C; [F (2, 18) = 0.2223, P=0.8028]), indicating that NMDAR conductance does not differentially contribute to spike generation in *Grin2a*-deficient pyramidal neurons under these conditions.

**Figure 2.**
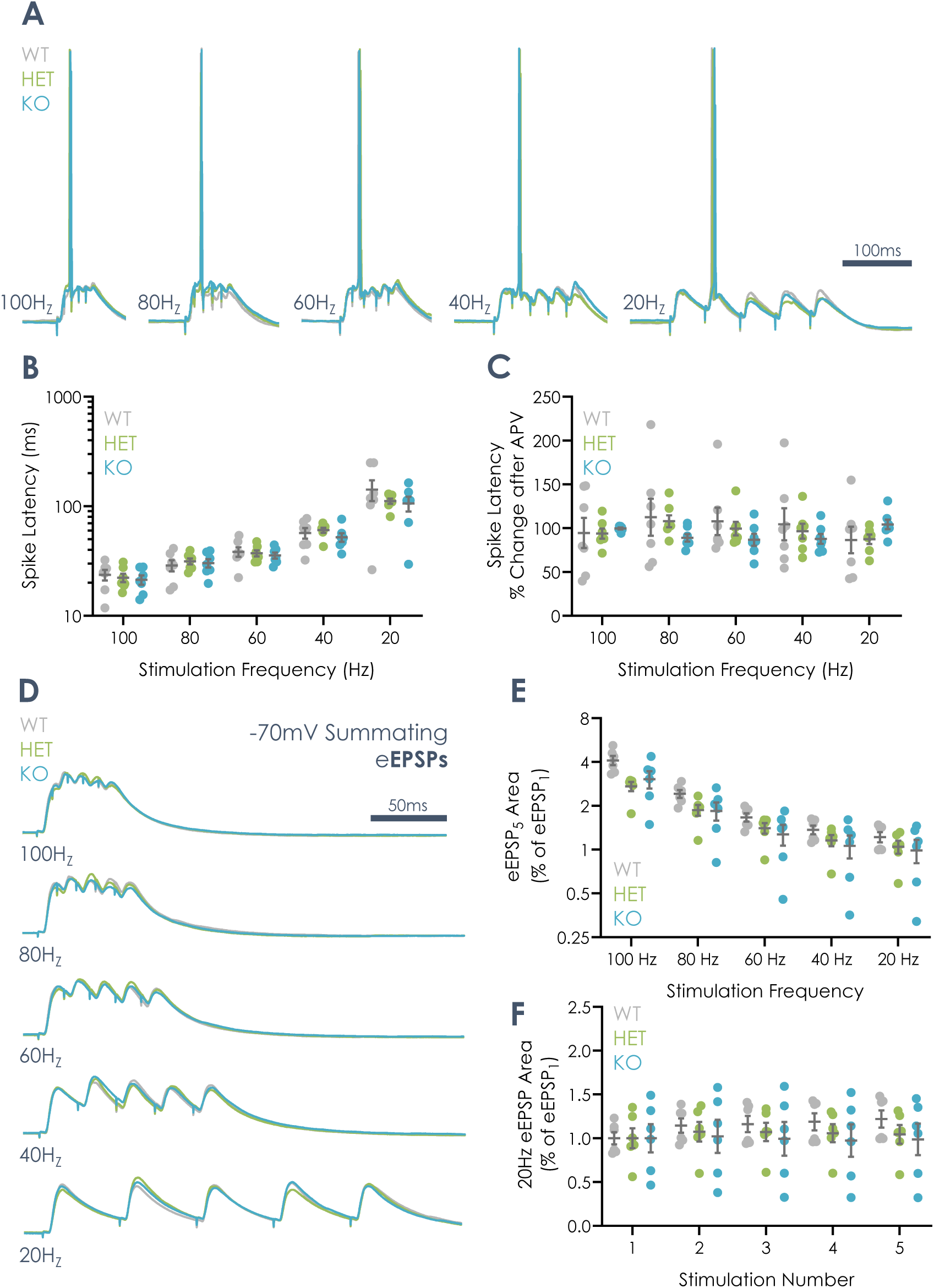
**Loss of *Grin2a* does not alter temporal summation of EPSPs or spike recruitment in CA1 pyramidal neurons**. (A) Example traces from CA1 pyramidal neurons from WT, HET and KO rats, showing responses to trains of 5 Schaffer collateral stimuli delivered at 100, 80, 60, 40 and 20 Hz. Membrane potential was held at −70 mV by injecting a bias current, and stimulus intensity was set to evoke a first subthreshold eEPSP of approximately 5-10 mV. (B) Spike latency (time from the first stimulus to action potential threshold) at each stimulation frequency revealed no effect of genotype (animal averages; N = 7 WT, 7 HET, 7 KO animals). (C) Change in spike latency (%) following wash-in of D-APV, at each stimulation frequency. There was no effect of genotype, indicating that NMDARs, including GluN2A-containing NMDARs, do not make a substantial contribution to synaptic integration and spike generation in pyramidal neurons under these conditions (animal averages; N = 7 WT, 7 HET, 7 KO animals). (D) As in (A), but with QX-314 included in the patch pipette to block action potential firing and reveal the underlying summating EPSP waveform. (E) Area of the 5th EPSP normalised to the area of the 1st EPSP, at each stimulation frequency. There was no effect of genotype (animal averages; N = 6 WT, 6 HET, 6 KO animals). (F) EPSP area for each of the 5 successive stimuli in a 20 Hz train. There was no effect of genotype at any stimulus position (animal averages; N = 6 WT, 6 HET, 6 KO animals). Data are mean ± SEM, with each point representing one animal. Statistical comparisons used ANOVA with Holm-Sidak post hoc tests. ns, not significant.

To examine subthreshold integration independently of action potentials, the protocol was repeated with intracellular QX −314. Normalised fifth/first EPSP area did not differ between genotypes at any stimulation frequency (Figure 2D-E; [F (2, 15) = 2.602, P=0.1071]). Analysis of individual EPSP areas across 20Hz trains likewise revealed no genotype effect at any stimulus position (Figure 2F; [F (2, 15) = 0.3197, P=0.7312]). These findings in an extended frequency rage further support earlier work from Yasmin *et al* (2025); providing robust evidence that *Grin2a* haploinsufficiency or knockout does not alter temporal summation of Schaffer collateral-evoked EPSPs in CA1 pyramidal neurons.

### GluN2A loss does not impair Hebbian plasticity at the Schaffer collateral synapse

Given the proposed role of GluN2A-containing NMDARs in LTP, we tested whether *Grin2a* loss altered Hebbian plasticity. HFS produced robust and sustained potentiation of eEPSC amplitude in WT, HET and KO neurons (Figure 3A-B). LTP magnitude at 30 min post-conditioning did not differ between genotypes (Figure 3C; [F (2, 19) = 0.5482, P=0.5868]). Thus, neither *Grin2a* haploinsufficiency nor complete knockout impairs HFS-induced LTP at Schaffer collateral synapses. Together with the preceding experiments, these findings indicate that CA1 pyramidal neuron basal transmission, integration and plasticity are largely preserved despite altered NMDAR deactivation kinetics.

**Figure 3.**
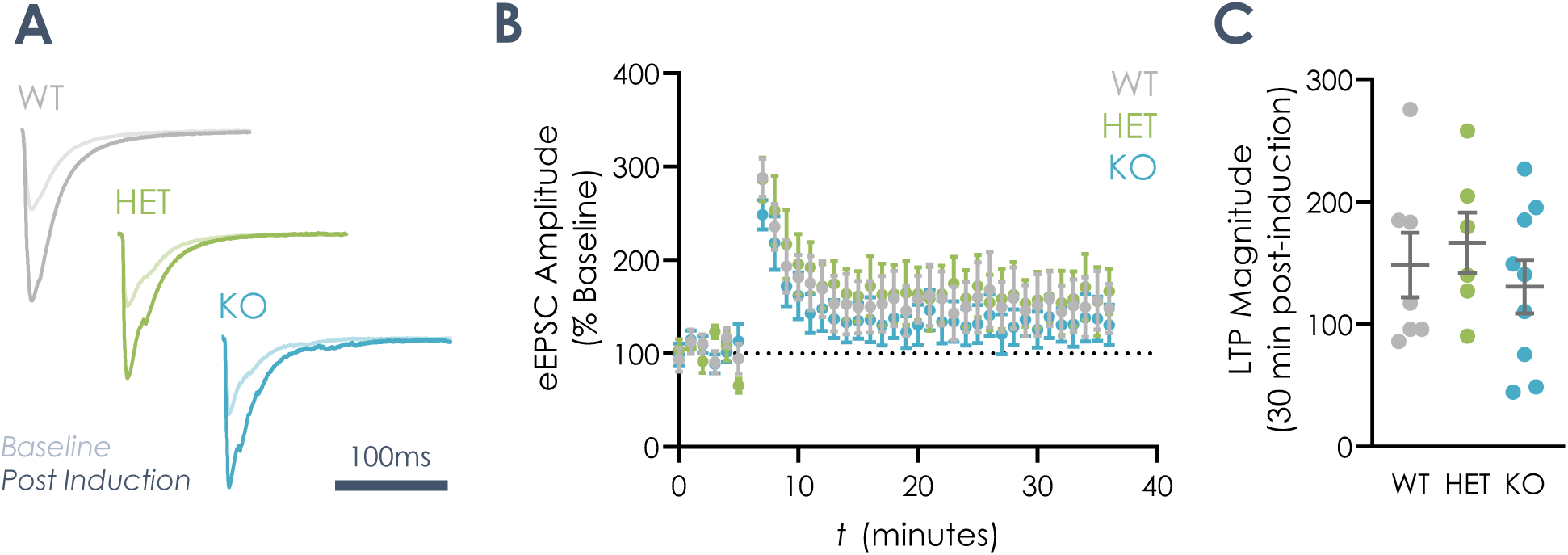
**Hebbian LTP at Schaffer collateral synapses onto CA1 pyramidal neurons is unaltered by Grin2a loss**. (A) Example eEPSC traces from CA1 pyramidal neurons from WT, HET and KO rats, recorded at baseline and 30 minutes post induction. Responses were evoked by stimulation of the Schaffer collaterals with a bipolar electrode, with cells voltage clamped at −70 mV. (B) Time course of eEPSC amplitude, normalised to baseline, for WT, HET and KO neurons. Recordings began with a 6-minute baseline, followed by conditioning and 30 minutes of post-induction recording. Conditioning consisted of two 1 s trains at 100 Hz, delivered 20 s apart. Potentiation was robust and sustained in all genotypes. (C) LTP magnitude, measured as eEPSC amplitude 30 minutes after conditioning as a percentage of baseline. There was no effect of genotype (animal averages; N = 7 WT, 6 HET, 9 KO animals). Data are mean ± SEM, with each point in (C) representing one animal. Statistical comparisons used one-way ANOVA with Holm-Sidak post hoc tests. ns, not significant.

### GluN2A loss selectively slows EPSP kinetics in OLM somatostatin interneurons but not parvalbumin interneurons

We next asked whether inhibitory interneuron subtypes are more sensitive to GluN2A loss. In parvalbumin interneurons, Schaffer collateral-evoked EPSP decay tau, halfwidth and area were unchanged across WT, HET and KO tissue (Figure 4D-G; [decay tau: F (2, 27) = 0.2486, P=0.7816; halfwidth: F (2, 27) = 0.08228, P=0.9212; area: F (2, 27) = 0.4145, P=0.6648]), indicating preserved EPSP waveform and integrated synaptic drive.

**Figure 4.**
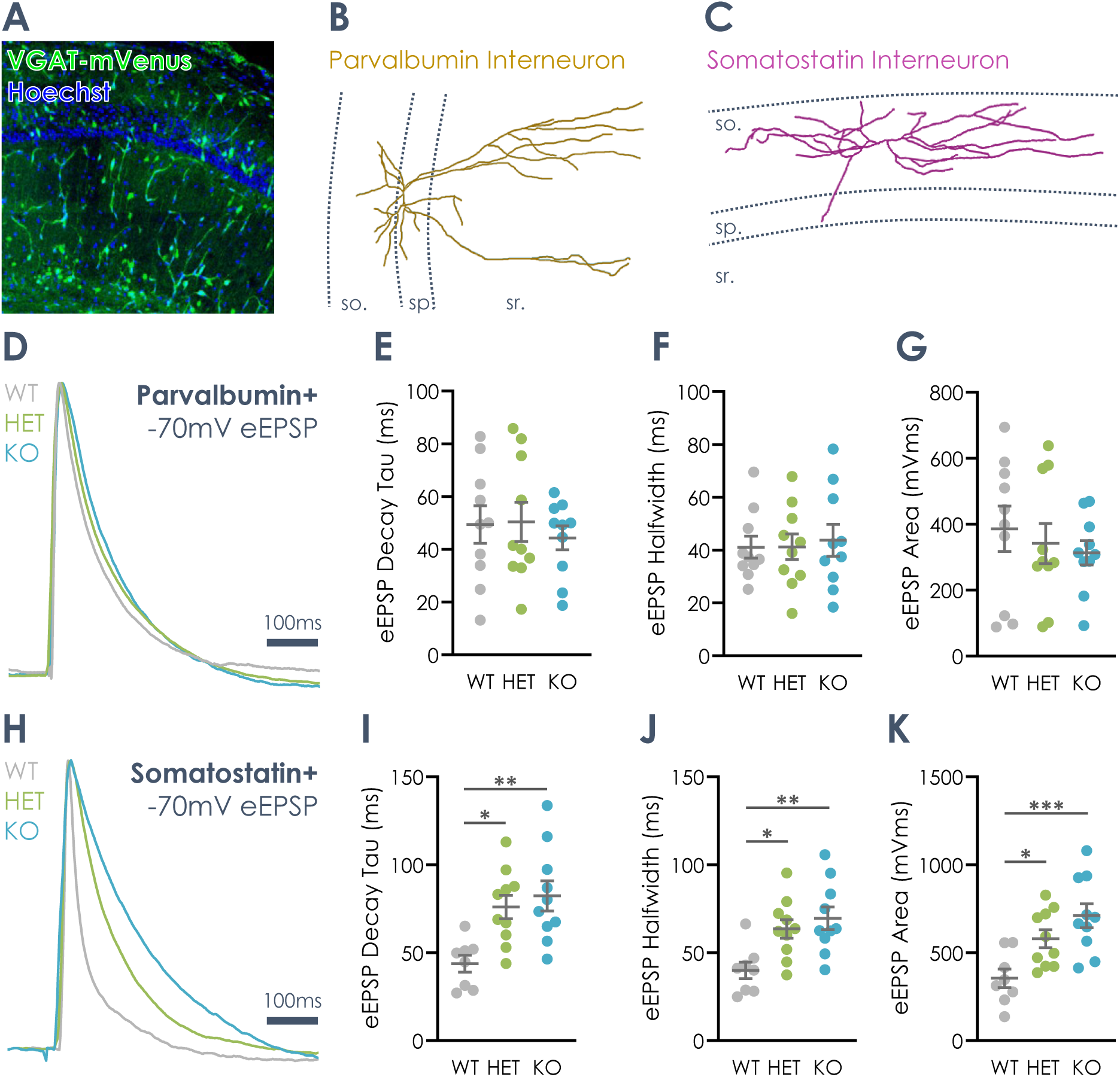
**Loss of GluN2A prolongs Schaffer collateral-evoked EPSPs in somatostatin interneurons but not parvalbumin interneurons**. (A) Example confocal image of the CA1 region of a VGAT-mVenus rat, showing mVenus fluorescence in hippocampal GABAergic interneurons. Interneurons were targeted for whole-cell recording by this fluorescence. (B) Reconstruction of a biocytin-filled parvalbumin (PV) interneuron, with its soma in stratum pyramidale. (C) Reconstruction of a biocytin-filled somatostatin (SST) interneuron, with its soma and dendrites in stratum oriens. (D) Example eEPSPs from PV interneurons from WT, HET and KO rats, evoked by Schaffer collateral stimulation with a bipolar electrode. Cells were held at −70 mV by injecting a bias current. (E-G) PV interneuron eEPSP decay τ (E), halfwidth (F) and area (G). None differed between genotypes (animal averages; N = 10 WT, 10 HET, 10 KO animals). (H) Example eEPSPs from SST interneurons from WT, HET and KO rats, recorded as in (D). (I–K) SST interneuron eEPSP decay τ (I), halfwidth (J) and area (K). All three measures were significantly greater in HET and KO tissue than in WT (animal averages; N = 8 WT, 10 HET, 10 KO animals). Data are mean ± SEM, with each point representing one animal. Statistical comparisons used one-way ANOVA with Holm-Sidak post hoc tests.

In contrast, somatostatin interneurons showed pronounced genotype-dependent EPSP broadening. eEPSP decay tau was significantly prolonged in HET and KO relative to WT, with corresponding increases in eEPSP halfwidth and area (Figure 4H-K; [decay tau: F (2, 25) = 7.736, P=0.0024; halfwidth: F (2, 25) = 7.061, P=0.0037; area: F (2, 25) = 8.781, P=0.0013]). Thus, GluN2A loss selectively prolongs excitatory synaptic potentials and increases depolarising charge transfer in somatostatin interneurons, consistent with a greater functional contribution of GluN2A-dependent NMDAR kinetics at these synapses.

### Reduced dendritic complexity and inhibitory input in somatostatin interneurons lacking GluN2A

To determine whether altered somatostatin eEPSP kinetics reflected broader changes in synaptic input or morphology, spontaneous synaptic currents and dendritic structure were examined. sEPSC amplitude, frequency and decay tau were unchanged across genotypes (Figure 5A-D; [amplitude: F (2, 22) = 0.9500, P=0.4020; frequency: F (2, 26) = 0.1769, P=0.8389; decay tau: F (2, 22) = 0.7106, P=0.5023]), indicating preserved spontaneous excitatory input onto somatostatin interneurons at −70mV.

**Figure 5.**
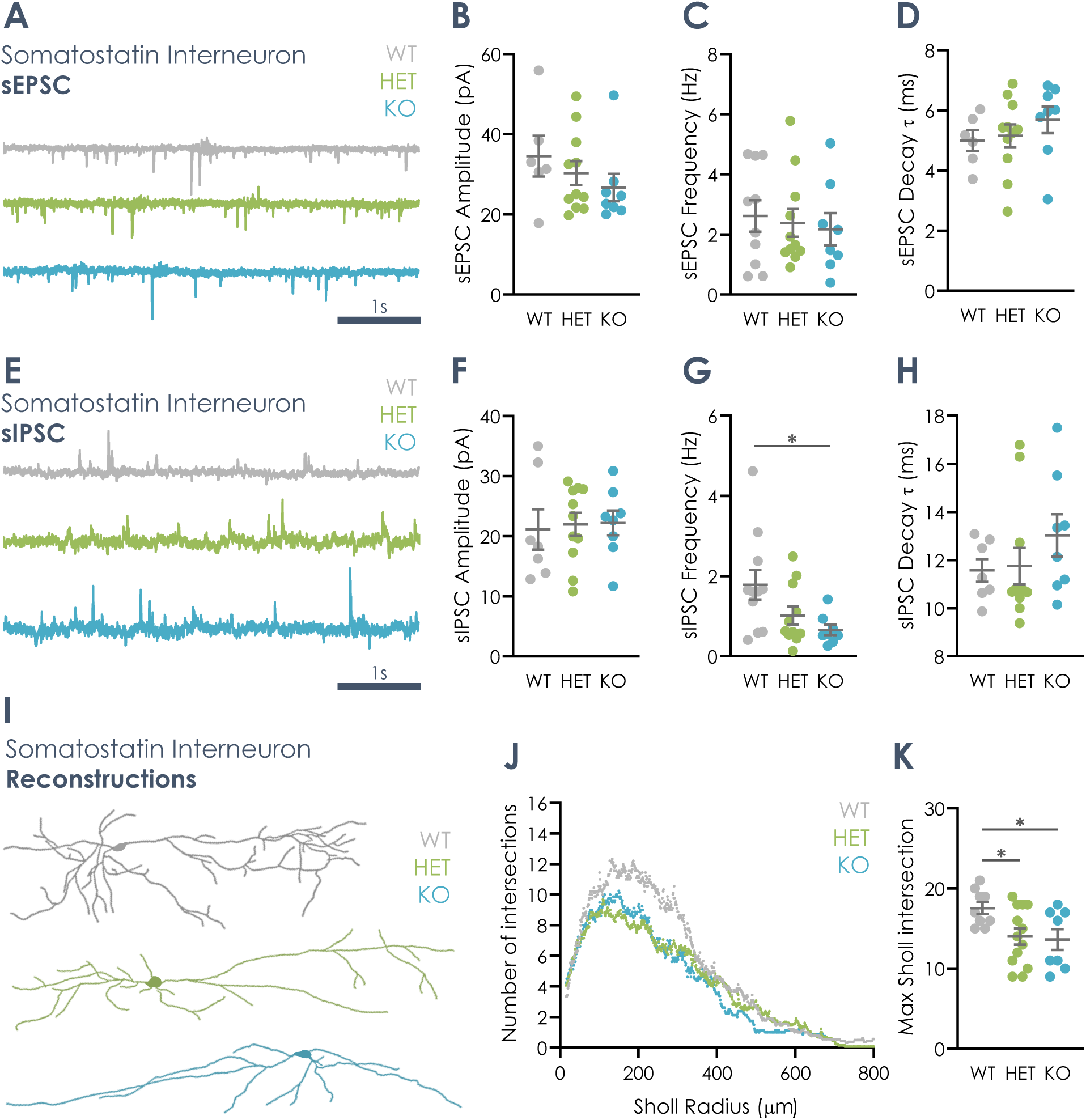
**Loss of GluN2A reduces spontaneous inhibitory input and dendritic complexity in SST interneurons without altering spontaneous excitatory input**. (A) Example sEPSC recordings from SST interneurons from WT, HET and KO rats, recorded in voltage clamp at −70 mV. (B-D) sEPSC amplitude (B), frequency (C) and decay τ (D). None differed between genotypes (animal averages; N = 6 WT, 11 HET, 8 KO animals). (E) Example sIPSC recordings from SST interneurons from WT, HET and KO rats, recorded in the same cells after changing the holding potential to 0 mV. (F-H) sIPSC amplitude (F), frequency (G) and decay τ (H). Amplitude and decay τ did not differ between genotypes, whereas sIPSC frequency was significantly reduced in HET and KO tissue (animal averages; N = 7 WT, 11 HET, 8 KO animals). (I) Example reconstructions of biocytin-filled SST interneurons from WT, HET and KO rats. (J) Sholl analysis, showing the number of dendritic intersections as a function of distance from the soma (Sholl radius, µm). (K) Maximum number of intersections from the Sholl analysis, which was significantly reduced in HET and KO cells compared with WT (animal averages; N = 9 WT, 13 HET, 8 KO animals). Data are mean ± SEM, with each point representing one animal. Statistical comparisons used one- way ANOVA with Holm-Sidak post hoc tests.

sIPSC amplitude and decay tau were also unchanged (Figure 5E, F, H; [amplitude: F (2, 23) = 0.04926, P=0.9520; decay tau: F (2, 23) = 1.020, P=0.3764]). However, sIPSC frequency was significantly reduced in KO tissue (Figure 5G; [F (2, 27) = 4.011, P=0.0298]), suggesting fewer functional inhibitory contacts or reduced activity of inhibitory afferents onto somatostatin interneurons.

Post-hoc morphological reconstruction revealed reduced dendritic complexity in *Grin2a*-deficient somatostatin interneurons. Sholl analysis showed fewer dendritic intersections across multiple radii in HET and KO cells, and maximum intersection number was significantly reduced relative to WT (Figure 5I-K; [F (2, 27) = 3.988, P=0.0304]). Thus, GluN2A loss is associated with simplified somatostatin interneuron dendritic morphology and reduced inhibitory input frequency, while spontaneous excitatory input remains intact.

### Intrinsic excitability and passive membrane properties across hippocampal cell populations

To determine whether synaptic phenotypes were accompanied by changes in intrinsic excitability, current-step responses were examined in parvalbumin and somatostatin interneurons. Action potential input-output relationships were unchanged across genotypes in parvalbumin interneurons; however, a subtle genotype effect was revealed in somatostatin interneurons such that cells from KO tissue showed greater firing at high levels of current injection (Figure 6A-D; [parvalbumin: F (2, 31) = 0.6311, P=0.5387; somatostatin: F (2, 30) = 3.543, P=0.0416]). Resting membrane potential, rheobase and membrane time constant were also unaffected in both cell types (Figure 6E-G; [parvalbumin resting membrane potential: F (2, 27) = 2.047, P=0.1487; parvalbumin rheobase: F (2, 27) = 1.012, P=0.3770; parvalbumin membrane time constant: F (2, 27) = 1.492, P=0.2429; somatostatin resting membrane potential: F (2, 34) = 0.02473, P=0.9756; somatostatin rheobase: F (2, 34) = 0.6178, P=0.5451; somatostatin membrane time constant: F (2, 34) = 1.002, P=0.3778]).

**Figure 6.**
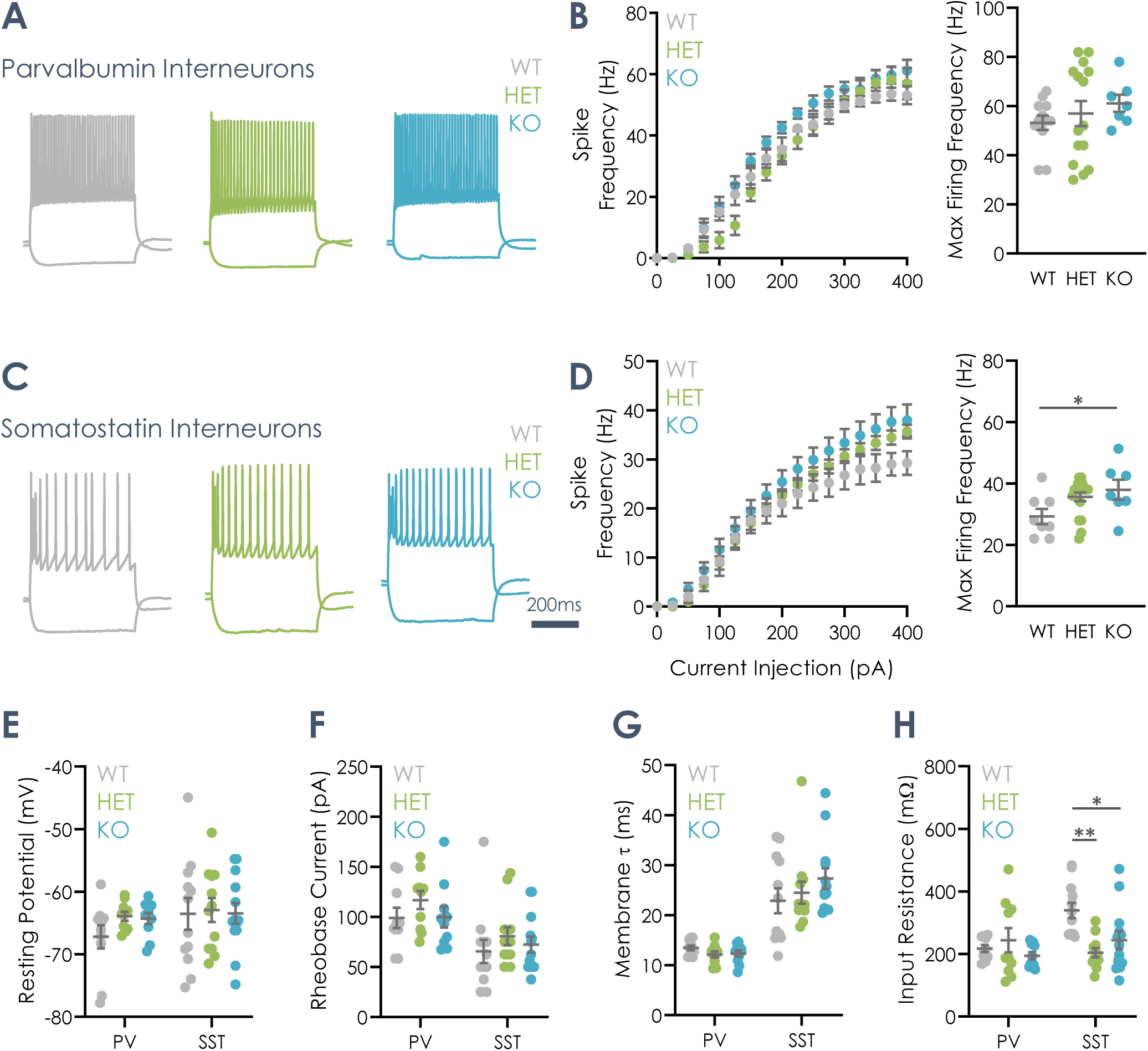
**GluN2A loss does not alter intrinsic excitability of PV interneurons, but reduces input resistance in SST interneurons**. (A) Example voltage responses of PV interneurons from WT, HET and KO rats to 500 ms current steps (−100 to +400 pA in 25 pA increments), recorded in current clamp from the apparent resting potential. (B) Input-output curve (left and max firing frequency (right) for PV interneurons (action potential frequency against injected current). There was no significant effect of genotype on max firing frequency (animal averages; N = 12 WT, 15 HET, 7 KO animals). (C) As in (A), for SST interneurons. (D) Input-output curve (left) and max firing frequency (right) for SST interneurons. There was a significant effect of genotype whereby cells in KO tissue showed a higher max firing frequency (animal averages; N = 8 WT, 18 HET, 7 KO animals). (E-H) Resting membrane potential (E), rheobase current (F), membrane time constant (G) and input resistance (H) for PV and SST interneurons from WT, HET and KO tissue. Resting membrane potential, rheobase and membrane time constant did not differ between genotypes in either cell type (PV animal averages; N = 10 WT, 10 HET, 10 KO animals and SST animal averages; N = 12 WT, 12 HET, 13 KO animals). Input resistance did not differ between genotypes in PV interneurons but was significantly lower in HET and KO SST interneurons than in WT (PV animal averages; N = 10 WT, 10 HET, 10 KO animals and SST animal averages; N = 12 WT, 12 HET, 13 KO animals). Data are mean ± SEM; for the summary plots (E–H) each point represents one animal. Statistical comparisons used ANOVA with Holm-Sidak post hoc tests.

Input resistance was unchanged in parvalbumin interneurons but significantly reduced in HET and KO somatostatin interneurons relative to WT (Figure 6H; [parvalbumin: F (2, 27) = 1.044, P=0.3658; somatostatin: F (2, 34) = 8.328, P=0.0011]). Despite this reduction, somatostatin firing output to somatic current injection is slightly hyperexcitable. These data indicate that GluN2A loss does not broadly alter intrinsic excitability in parvalbumin interneurons, but reveals modest changes to the intrinsic excitability of somatostatin interneurons. As such, these data indicate that somatostatin interneurons appear functionally sensitive to loss of GluN2A at both synaptic, structural and cell intrinsic functional scales.

### Amplified synaptic summation and activity-dependent firing in OLM somatostatin interneurons lacking GluN2A

Because somatostatin interneuron EPSPs were prolonged, we tested whether repetitive Schaffer collateral input produced enhanced temporal summation. In parvalbumin interneurons, normalised fifth/first EPSP area was reduced by GluN2A loss at high stimulation frequencies, which may reflect an immature electrophysiological profile or altered network state promoting increased inhibitory drive onto these interneurons (Figure 7A, C; [F (2, 22) = 13.74, P=0.0001]). However, individual EPSP areas during 20Hz trains were unchanged (Figure 7E; [F (2, 22) = 0.4284, P=0.6568]). Thus, parvalbumin interneuron synaptic integration is modestly reduced by *Grin2a* loss under high frequency stimulation (>60Hz) that is not anticipated to arise directly from eEPSP decay kinetics.

**Figure 7.**
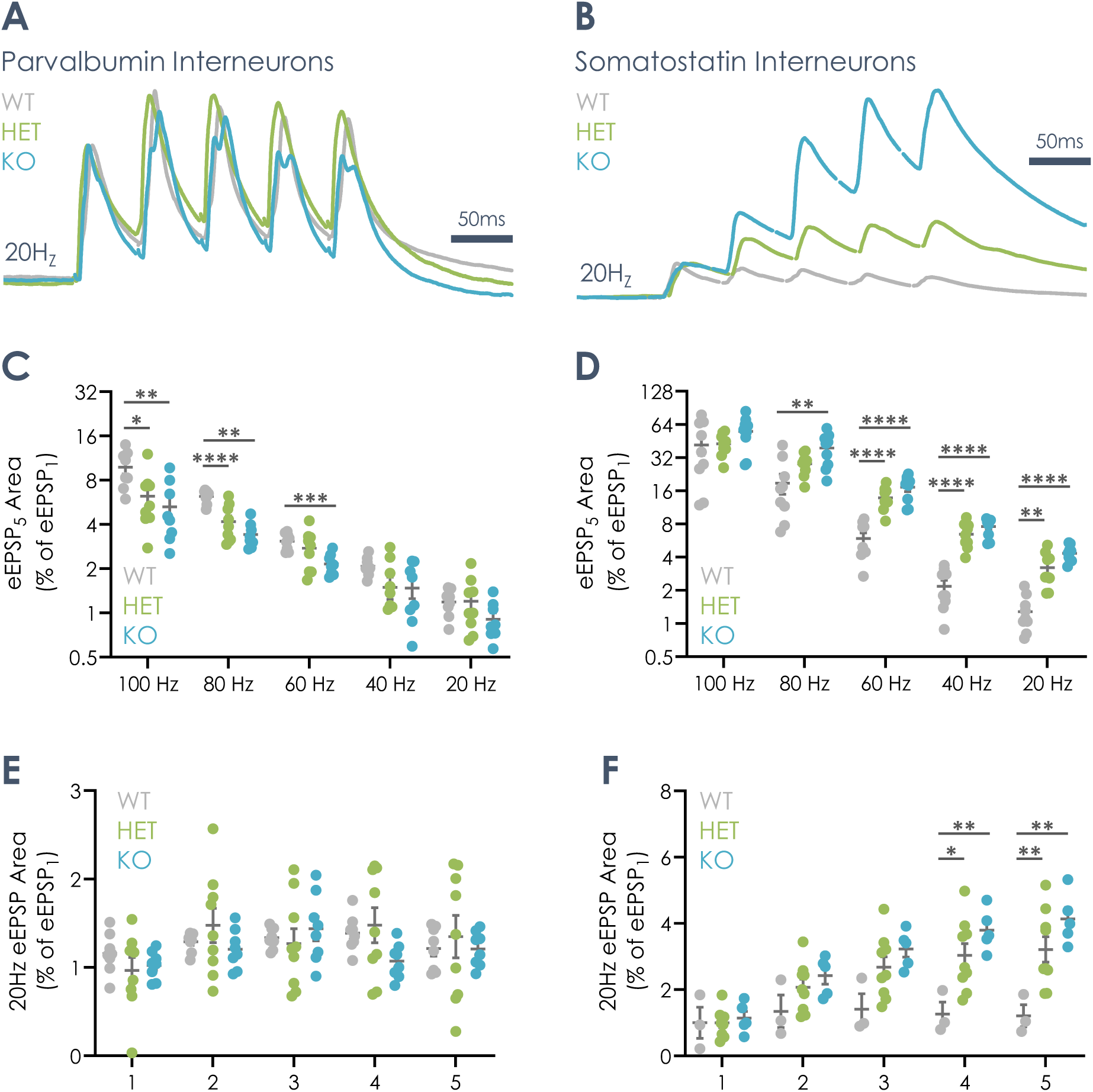
Loss of GluN2A enhances temporal summation of Schaffer collateral-evoked EPSPs in SST interneurons. (A) Example traces from CA1 PV interneurons from WT, HET and KO rats, showing responses to a 20 Hz train of 5 Schaffer collateral stimuli. Recordings were made in current clamp with QX-314 in the pipette to block action potential firing and reveal the underlying summating EPSP waveform. Membrane potential was held at −70 mV by injecting a bias current. (B) As in (A), for CA1 SST interneurons. Summating EPSPs have a visibly larger amplitude in HET and KO cells. (C) Area of the 5th EPSP normalised to the area of the 1st EPSP, at 100, 80, 60, 40 and 20 Hz, in PV interneurons. There was no effect of genotype (animal averages; N = 8 WT, 9 HET, 8 KO animals). (D) As in (C), for SST interneurons. There was a significant effect of genotype, with area greater in HET and KO tissue, most pronounced at lower stimulation frequencies (animal averages; N = 9 WT, 9 HET, 9 KO animals). (E) EPSP area for each of the 5 successive stimuli in a 20 Hz train in PV interneurons. There was no effect of genotype (animal averages; N = 8 WT, 9 HET, 8 KO animals). (F) As in (E), for SST interneurons. There was a significant effect of genotype, with EPSP area greater in HET and KO tissue, most pronounced at later stimuli in the train (animal averages; N = 9 WT, 9 HET, 9 KO animals). Data are mean ± SEM, with each animal contributing a single averaged value. Statistical comparisons used ANOVA with Holm-Sidak post hoc tests.

In somatostatin interneurons, however, repetitive stimulation produced markedly greater EPSP summation in HET and KO tissue (Figure 7B). Normalised fifth EPSP area was significantly increased in *Grin2a*-deficient somatostatin interneurons, with the largest genotype effects at lower stimulation frequencies, particularly 20 and 40Hz (Figure 7B, D; [F (2, 24) = 9.037, P=0.0012]). This frequency dependence is consistent with prolonged EPSP decay increasing residual depolarisation between more widely spaced stimuli. Analysis of individual EPSPs during 20Hz trains showed that genotype differences emerged from the second stimulus onward and increased across the train, reaching a maximum at the fifth stimulus (Figure 7F; [F (2, 24) = 9.037, P=0.0012]). Thus, the prolonged EPSP kinetics caused by GluN2A loss translate into progressive, frequency-dependent amplification of synaptic integration in somatostatin interneurons.

Finally, we tested whether enhanced somatostatin summation increased action potential output. Parvalbumin interneuron spike counts during 100-20Hz five-stimulus trains were unchanged across genotypes (Figure 8A-B; [F (2, 22) = 1.485, P=0.2483]), consistent with preserved synaptic integration. In somatostatin interneurons, HET and KO cells fired more action potentials than WT during repetitive Schaffer collateral stimulation (Figure 8C-D; [F (2, 22) = 22.80, P<0.0001]). This increase was largest at lower frequencies, particularly below 60Hz, paralleling the frequency dependence of enhanced EPSP summation.

**Figure 8.**
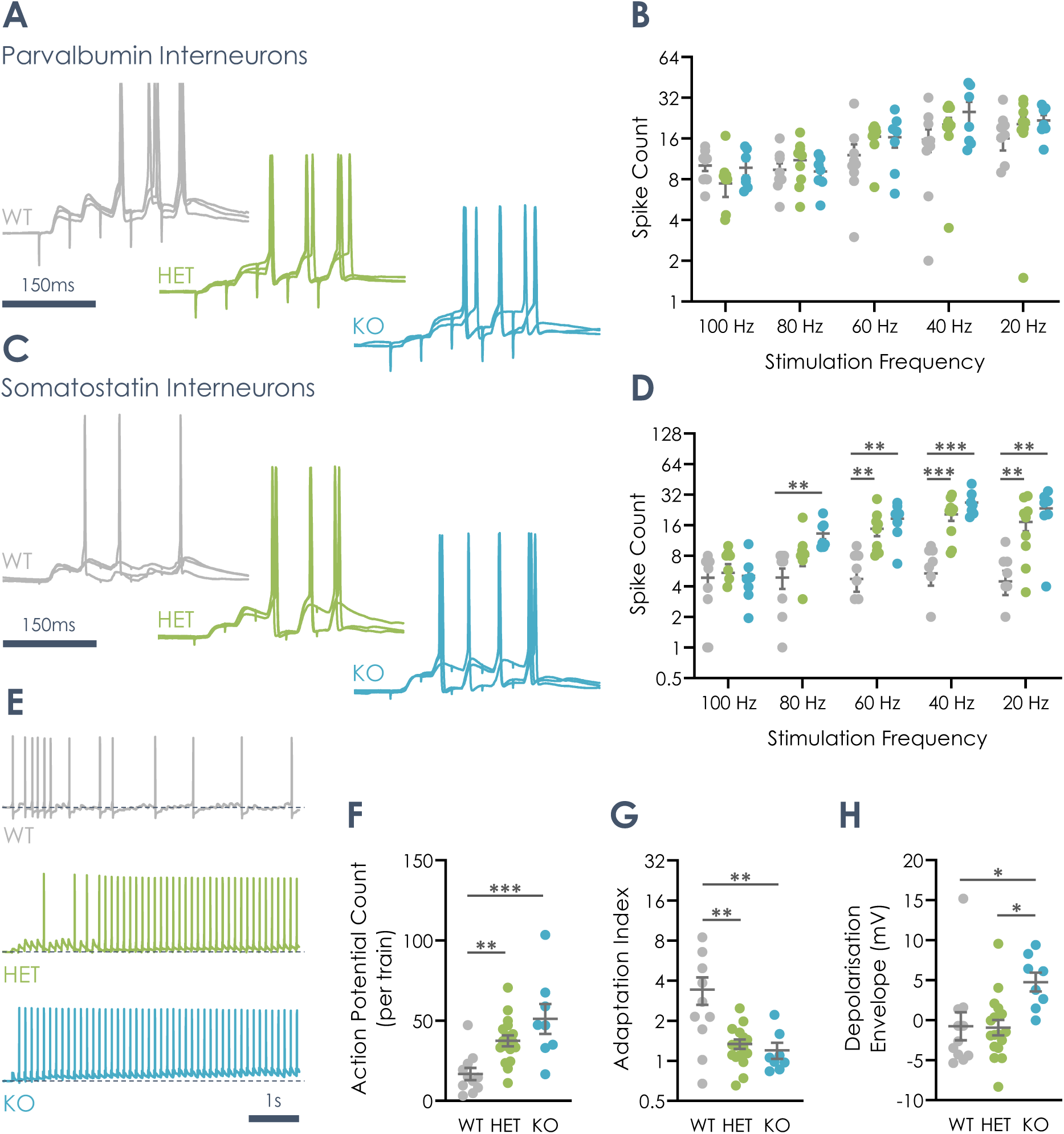
**Loss of GluN2A enhances synaptically evoked spiking in SST interneurons, but not PV interneurons**. (A) Example traces from CA1 PV interneurons from WT, HET and KO rats, showing responses to a 20 Hz train of 5 Schaffer collateral stimuli. Recordings were made in current clamp without QX-314 in the pipette, so action potential firing was intact. Membrane potential was held at − 70 mV by injecting a bias current. (B) Spike count per train in PV interneurons at 100, 80, 60, 40 and 20 Hz. There was no effect of genotype (animal averages; N = 9 WT, 9 HET, 7 KO animals). (C) As in (A), for CA1 SST interneurons. (D) As in (B), for SST interneurons. Spike count was significantly greater in HET and KO cells, most markedly at lower stimulation frequencies (animal averages; N = 9 WT, 9 HET, 7 KO animals). (E) Example traces from CA1 SST interneurons from WT, HET and KO rats, showing responses to 6 s of 8 Hz (theta-frequency) Schaffer collateral stimulation. (F) Number of action potentials per 8 Hz train, which was significantly greater in HET and KO SST interneurons than in WT (animal averages; N = 11 WT, 17 HET, 8 KO animals). (G) Spike-frequency adaptation across the 8 Hz train, which was reduced in HET and KO SST interneurons (animal averages; N = 11 WT, 17 HET, 8 KO animals). (H) Depolarisation envelope, the slow cumulative depolarisation that builds across the 8 Hz train, measured relative to the pre-train baseline (animal averages; N = 11 WT, 17 HET, 8 KO animals). It was significantly greater in KO SST interneurons. Data are mean ± SEM, with each animal contributing a single averaged value. Statistical comparisons used ANOVA with Holm-Sidak post hoc tests.

To assess sustained theta-range recruitment, somatostatin interneurons were stimulated at 8Hz for 6 seconds. HET and KO somatostatin interneurons generated significantly more action potentials than WT (Figure 8E-F; [F (2, 33) = 9.986, P=0.0004]) and showed reduced spike frequency adaptation (Figure 8G; [F (2, 33) = 10.34, P=0.0003]), maintaining elevated firing throughout the train. KO cells also exhibited a significantly larger depolarisation envelope (Figure 8H; [F (2, 33) = 4.832, P=0.0144]), consistent with greater accumulation of slow NMDAR-mediated depolarisation during sustained input. Together, these data show that GluN2A loss selectively enhances somatostatin interneuron synaptic integration and spike output, while leaving parvalbumin interneurons and CA1 pyramidal neurons largely unaffected. This OLM somatostatin interneuron hyperactivation provides a candidate circuit mechanism through which *GRIN2A* haploinsufficiency may disrupt hippocampal network dynamics.

## Discussion

In this study, we identify a selective and cell-type-specific consequence of GluN2A loss within hippocampal CA1 circuitry. Despite clear evidence that *Grin2a* haploinsufficiency and knockout slow NMDAR-mediated current deactivation under voltage clamp, basal excitatory transmission, temporal integration and HFS-induced LTP at Schaffer collateral synapses onto CA1 pyramidal neurons were largely preserved. Parvalbumin interneurons were similarly unaffected across measures of EPSP kinetics and intrinsic excitability, while temporal summation appears modestly impaired by GluN2A loss in this cell type. In contrast, OLM somatostatin interneurons displayed a pronounced gain-of-function phenotype: Schaffer collateral-evoked EPSPs were broadened, EPSP area was increased, temporal summation was enhanced, and repetitive synaptic stimulation drove substantially greater action potential output. These effects were accompanied by reduced dendritic complexity, reduced spontaneous inhibitory input frequency and intrinsic hyperexcitability at high levels of current injection. Together, our data indicate that GluN2A loss does not produce a generalised impairment of CA1 excitatory synaptic function. Instead, GluN2A loss appears to reveal distinct alterations in a cell type-dependent manner; most notably enhancing excitatory synaptic integration in OLM somatostatin interneurons, identifying this cell type as a selective locus of vulnerability to *GRIN2A* disruption.

### OLM somatostatin interneurons are selectively sensitive to GluN2A loss

GluN2A-containing NMDARs are classically associated with faster synaptic NMDAR kinetics, and their developmental incorporation into synapses is thought to sharpen the temporal window for coincidence detection and plasticity (Paoletti et al., 2013). Consistent with this, voltage-clamp recordings from CA1 pyramidal neurons revealed a gene-dose-dependent slowing of NMDAR-mediated eEPSC decay in HET and KO tissue. This confirms that GluN2A loss alters synaptic NMDAR kinetics and is consistent with compensatory enrichment of slower GluN2B-containing receptors (Elmasri et al., 2022). However, this kinetic change did not translate into detectable changes in pyramidal neuron eEPSP waveform, AMPA/NMDAR ratio, temporal summation, spike latency or HFS-induced LTP.

This apparent dissociation between altered NMDAR current kinetics and preserved pyramidal neuron function suggests that, at least under the stimulation and recording conditions used here, the pyramidal cell synaptic GluN2A-NMDAR pool is not rate-limiting for basal EPSP integration or conventional Schaffer collateral LTP. Several factors may contribute to this resilience. At resting membrane potentials, Mg²⁺ block limits NMDAR contribution to individual EPSPs, making AMPAR-mediated depolarisation the dominant determinant of spike recruitment during brief Schaffer collateral trains. In addition, compensatory redistribution of GluN2B-containing NMDARs may preserve total NMDAR charge transfer and support sufficient Ca²⁺ entry for LTP induction. Finally, the HFS protocol used here may provide a strong induction stimulus that overcomes subtle differences in NMDAR subunit composition. Thus, rather than abolishing pyramidal neuron NMDAR function, GluN2A loss appears to alter receptor kinetics in a manner that is measurable under voltage clamp but functionally buffered in pyramidal neurons during the assays performed.

The absence of a phenotype at the level of cellular output in parvalbumin interneurons further argues against a widespread disruption of excitatory transmission across hippocampal inhibitory circuits. Parvalbumin cells displayed normal EPSP kinetics, intrinsic excitability and synaptically driven firing, albeit with a modest reduction in summation at high frequencies. This is notable because parvalbumin interneurons are frequently implicated in epilepsy and in *GRIN2A*-related neurodevelopmental dysfunction (Camp et al., 2023; Cardis et al., 2018; Hosseini et al., 2025). Our findings do not exclude parvalbumin interneuron involvement in disease progression, but suggest that, in CA1, parvalbumin cell dysfunction may not arise from a primary alteration in their Schaffer collateral-evoked excitatory synaptic integration. Instead, parvalbumin phenotypes reported in *Grin2a*-deficient models may emerge indirectly, potentially through altered developmental interactions with other interneuron populations, including OLM somatostatin cells (Mòdol et al., 2024).

The central finding of this study is that GluN2A loss selectively prolongs excitatory synaptic potentials in OLM somatostatin interneurons and converts this kinetic change into enhanced spike output. The increase in EPSP decay tau, halfwidth and area indicates that each excitatory event delivers a longer-lasting depolarising drive in HET and KO cells. During repetitive stimulation, this prolonged depolarisation produces greater residual voltage at the time of subsequent inputs, thereby amplifying temporal summation. The frequency dependence of this effect provides important mechanistic support: genotype differences were greatest at 20-40Hz, where WT EPSPs have sufficient time to decay between stimuli, and were attenuated at higher frequencies, where EPSP overlap is already substantial in all genotypes. This relationship strongly supports the interpretation that prolonged EPSP kinetics are the primary driver of enhanced OLM somatostatin integration.

The expected underpinning mechanism is a shift in NMDAR composition away from GluN2A-containing receptors and toward slower GluN2B-dominated receptors. This is supported by the prior demonstration that eEPSCs onto somatostatin interneurons display a significantly increased sensitivity to ifenprodil - a GluN2B-selective antagonist - after GluN2A deletion; where parvalbumin and cholecystokinin interneurons do not (Booker et al., 2021). GluN2B-containing NMDARs deactivate more slowly than GluN2A-containing receptors and would be expected to prolong synaptic conductance, increase charge transfer, and favour temporal summation over tens to hundreds of milliseconds (Monyer et al., 1994; Vicini et al., 1998). Why this effect is functionally segregated to OLM somatostatin interneurons, however, remains an open question. One possibility is that OLM somatostatin synapses normally contain a higher relative proportion of GluN2A-containing NMDARs, such that GluN2A loss produces a larger fractional change in receptor kinetics (Lalanne et al., 2016; Wang and Gao, 2009). Differences in dendritic architecture and electrotonic filtering may also be important. OLM somatostatin cells possess horizontally oriented dendrites in stratum oriens and receive excitatory input onto dendritic compartments that may favour the influence of slow synaptic conductances on somatic voltage (Morabito et al., 2025). In this context, a prolongation of synaptic NMDAR current could be amplified by dendritic filtering into a measurable broadening of the somatic EPSP.

### Enhanced somatostatin recruitment has important circuit implications

OLM somatostatin interneurons occupy a distinctive position in CA1 microcircuits. Their axons target the distal apical dendrites of pyramidal neurons in stratum lacunosum-moleculare, where they regulate entorhinal cortical input, while their activity can also influence local disinhibitory motifs involving other interneuron classes (Fernández-Arroyo et al., 2025; Liguz-Lecznar et al., 2022). By controlling dendritic inhibition, OLM somatostatin cells help determine the balance between CA3-driven Schaffer collateral input and temporoammonic input from entorhinal cortex (Leão et al., 2012). Hyperactivation of OLM somatostatin interneurons following GluN2A loss could therefore bias information flow through CA1 by altering the relative weighting of these convergent pathways, reducing excitatory cortical drive through the temporoammonic pathway.

Such a shift may have consequences for hippocampal computations that depend on the integration of internally generated and sensory-contextual representations. CA3 input is commonly associated with pattern completion and retrieval of stored representations, whereas entorhinal input conveys cortical contextual and spatial information important for updating and encoding (Cabral et al., 2014; Colgin et al., 2009; Guardamagna et al., 2025). Excessive OLM somatostatin interneuron-mediated dendritic inhibition could suppress distal dendritic excitability occluding integration of temporoammonic input, potentially impairing spatial learning, place field stability or context-dependent memory encoding (Abbas et al., 2018; Honoré and Lacaille, 2022). This possibility is consistent with evidence that somatostatin interneurons contribute to place field formation and with reports of spatial learning impairments in *Grin2a*-deficient mice (Acutain et al., 2021; Hosseini et al., 2025; Udakis et al., 2025).

The enhanced firing of OLM somatostatin interneurons during theta-frequency stimulation is particularly relevant. OLM cells are strongly recruited during hippocampal theta rhythms and are positioned to regulate dendritic excitability across behavioural states associated with exploration and memory encoding (Antonoudiou et al., 2020; Brockway et al., 2023; Huang et al., 2020). Our finding that HET and KO OLM somatostatin cells fire more during sustained 8Hz stimulation, show reduced spike frequency adaptation, and in KO tissue develop a larger depolarisation envelope suggests that GluN2A loss may increase OLM gain during theta-like activity. This could reshape theta-phase-dependent dendritic inhibition and alter the timing of pyramidal cell input integration. Given the importance of precisely timed inhibition for hippocampal oscillations, altered somatostatin interneuron functional output at this frequency may have broad consequences for network synchrony and information processing.

### GluN2A deletion drives dendritic and inhibitory synapse alterations in OLM somatostatin interneurons

In addition to altered excitatory integration, *Grin2a*-deficient OLM somatostatin interneurons showed reduced dendritic complexity and sIPSC frequency. The reduction in dendritic arborisation suggests that GluN2A contributes to the development or maintenance of somatostatin interneuron structure. NMDAR-dependent expression is known to regulate dendritic growth, stabilisation and synapse maturation in excitatory cells, as such, the loss of GluN2A may similarly perturb these processes during development of inhibitory interneurons such as somatostatin OLM cells (Acutain and Baez, 2024; Chu et al., 2026; Kannangara et al., 2014; Yasmin et al., 2025). The fact that dendritic simplification was evident in both HET and KO tissue, with no clear additional reduction in KO cells, suggests that partial GluN2A loss may be sufficient to disrupt this structural programme. Because the morphological analysis is representational of a P21-34 developmental window, it remains unclear whether dendritic simplification is a developmental consequence of GluN2A loss or a compensatory response to altered activity. Longitudinal developmental analysis will be required to distinguish these possibilities, and shed further light on the impact of GluN2A loss on somatostatin interneuron development and function.

The reduced frequency of spontaneous inhibitory input, in the absence of changes in sIPSC amplitude or decay, points to a decrease in the number of functional inhibitory contacts onto OLM somatostatin cells - which may be scaled down with reduced dendritic arborisation - or may reflect reduced spontaneous activity of presynaptic inhibitory afferents. Functionally, reduced inhibition would be expected to further favour OLM recruitment in vivo, complementing the enhanced excitatory summation and cellular hyperexcitability observed during evoked stimulation and current injection, respectively. However, because spontaneous recordings cannot distinguish changes in synapse number from changes in presynaptic activity, additional approaches will be required to define the source of this inhibitory deficit.

The reduction in input resistance in HET and KO OLM somatostatin interneurons is also notable. Lower input resistance would normally reduce voltage responses to synaptic current and oppose hyperexcitability. The fact that EPSP summation and spike output were nevertheless increased indicates that the prolongation of synaptic conductance is sufficiently strong to overcome this passive dampening. The reduced input resistance may reflect altered membrane conductance or homeostatic compensation in response to the reduced inhibitory drive.

### Relationship to plasticity and prior GluN2A literature

The preservation of HFS-induced LTP in CA1 pyramidal neurons may appear surprising given the common association between GluN2A-containing NMDARs and LTP (Li et al., 2007; Liu et al., 2004). However, previous work has shown that the relationship between GluN2A and LTP is not absolute and can heavily depend on developmental stage, induction protocol, synapse type and experimental preparation (Gerkin et al., 2007; Kohl et al., 2011; Köhr et al., 2003; Zhang et al., 2008). The present data support a model in which GluN2A shapes NMDAR kinetics rather than acting as an obligatory gate for potentiation (Erreger et al., 2005; Shipton and Paulsen, 2014). Under strong HFS conditions at Schaffer collateral synapses, residual NMDAR signalling in HET and KO pyramidal neurons appears sufficient to support robust LTP. However, this does not exclude the possibility of a more subtle plasticity phenotype under other conditions: GluN2A loss may alter the threshold, timing dependence or input specificity of plasticity without changing the magnitude of HFS-induced LTP. Protocols that recruit weaker or more physiological patterns of activity, spike-timing-dependent plasticity, theta-burst stimulation, or plasticity at temporoammonic inputs may reveal deficits not captured here. In addition, OLM somatostatin hyperactivation could influence pyramidal neuron plasticity indirectly in intact circuits by increasing dendritic inhibition during induction. It has previously been shown that somatostatin interneuron hyperactivation can increase Schaffer collateral LTP through a disinhibition of local parvalbumin interneuron inputs in the stratum radiatum (Leão et al., 2012). Thus, we cannot rule out opposing impacts of GluN2A loss, such that, direct impacts at the Schaffer collateral synapse - reducing LTP - may be offset by a somatostatin interneuron-dependent enhancement of plasticity at this input. Consequently, the absence of a definitive pyramidal neuron LTP deficit in this study should be interpreted as evidence that canonical Schaffer collateral HFS-LTP is intact in this germline developmental model, rather than as evidence that all forms of hippocampal plasticity are unaffected by GluN2A loss.

### Implications for *GRIN2A*-related neurodevelopmental disorders

*GRIN2A* loss-of-function variants are strongly associated with epilepsy-aphasia spectrum disorders, including self-limited epilepsy with centrotemporal spikes, Landau-Kleffner syndrome and epileptic encephalopathy with continuous spike-and-wave during sleep (Strehlow et al., 2019). Yet, the mechanisms linking *GRIN2A* haploinsufficiency to seizures and language dysfunction remain unresolved. Our findings suggest that the relevant circuit defect may not be a simple global alteration in excitatory drive; instead, GluN2A loss can produce a selective gain-of-function in a defined inhibitory interneuron subpopulation.

At first glance, enhanced firing of inhibitory interneurons may seem inconsistent with epileptiform network hyperexcitability. However, interneuron dysfunction can promote epilepsy in multiple ways depending on subtype, timing and circuit context. Excessive OLM activity could alter disinhibitory control of parvalbumin interneurons, and distort the balance of dendritic versus perisomatic inhibition (Cottam et al., 2013; Liguz-Lecznar et al., 2022; Xu et al., 2013). Moreover, if OLM hyperactivity occurs during development, it could perturb maturation of other interneuron populations. This is particularly relevant given evidenc e that parvalbumin interneuron maturation depends on somatostatin interneuron activity and that GluN2A loss delays parvalbumin interneuron electrophysiological maturation (Camp et al., 2023; Mòdol et al., 2024). Our data provide a plausible upstream mechanism: GluN2A loss selectively enhances excitatory drive onto OLM somatostatin interneurons, which could alter somatostatin interneuron signalling throughout development and impair subsequent maturation parvalbumin interneuron electrophysiological phenotypes. It will therefore be important to determine when OLM somatostatin EPSP prolongation, dendritic simplification and inhibitory input loss first emerge, whether they precede parvalbumin maturation delay, and whether they persist or compensate in adulthood.

### Limitations and future directions

Although this study examined pyramidal neurons, parvalbumin interneurons and OLM somatostatin interneurons, CA1 contains many additional interneuron subtypes. Other populations, including cholecystokinin-positive basket cells, neurogliaform cells, bistratified cells and vasoactive intestinal peptide interneurons, may also be sensitive to GluN2A loss. A complete circuit-level understanding will require broader interneuron subtype profiling.

Moreover, the experiments described here were performed in acute slices using controlled Schaffer collateral stimulation. This approach provides strong mechanistic resolution but cannot fully capture the dynamics of intact hippocampal networks, including neuromodulation, long-range projection inputs, behavioural state dependence and sleep-associated shifts in network dynamics. Future work using *in vivo* recordings coupled with optogenetic cell-specific modulation will be crucial to link *Grin2a*-mediated dysfunction of somatostatin interneuron physiology to behavioural output. Given the known association between *Grin2a* loss and robust phenotypes in mouse models, future studies should examine spatial memory, place field formation, vocalisation and seizure propensity. Linking the cellular phenotype identified here to behavioural and network outputs will be essential for understanding how *GRIN2A* haploinsufficiency drives pathophysiology.

## Conclusion

Our findings reveal that GluN2A loss has a highly selective impact on CA1 hippocampal circuitry. Rather than causing broad impairment of pyramidal neuron synaptic transmission or plasticity, *Grin2a* haploinsufficiency and knockout preferentially prolong excitatory synaptic potentials in OLM somatostatin interneurons. This synaptic slowing amplifies temporal summation, increases theta-range and train-evoked firing, and is accompanied by reduced dendritic complexity and reduced inhibitory input. These data identify OLM somatostatin interneurons as a primary functional locus of GluN2A-dependent regulation in CA1 and suggest that their hyperactivation may contribute to hippocampal circuit dysfunction in *GRIN2A*-related epilepsy-aphasia disorders.

## Acknowledgements

This work was funded by the European Union’s Horizon 2020 research and innovation programme under the Marie Sklodowska- Curie grant agreement No. 101202271 and the Simons Initiative for the Developing Brain. We thank the team at the University of Edinburgh IMPACT Imaging Facility and the BVS animal facility staff for their support in this project.

## Author Contributions

**Daniel Hunter**: Conceptualisation; Data curation; Formal analysis; Validation; Investigation; Visualisation; Writing - original draft; Writing - review and editing. **Hannah Kaese**: Data curation; Formal analysis; Validation; Investigation. **Peter Kind**: Supervision; Funding acquisition. **David Wyllie**: Conceptualisation; Data curation; Supervision; Funding acquisition; Visualisation; Project administration; Writing - review and editing.

